# A poly(A) polymerase and oligo(dT)-dependent method for the purification of engineering circular RNAs that enhances protein expression in eukaryotic cells

**DOI:** 10.1101/2024.12.27.630476

**Authors:** Lulu Wei, Mengjuan Zhang, Haifeng Zhang, Haiyan Zhao, Yingwu Mei

**Author notes:** Corresponding Author: Yingwu Mei, Department of Biochemistry and Molecular Biology, School of Basic Medical Sciences, Zhengzhou University, Zhengzhou, Henan, 450001, China.

## Abstract

Circular RNA (circRNA) has garnered significant attention as a promising therapeutic tool due to its remarkable stability and resistance to exonuclease degradation. However, the development of scalable and efficient purification methods remains a critical challenge. In this study, we propose and evaluate a novel purification strategy for engineered circular RNA (PAPdt), which leverages poly(A) polymerase and oligo(dT) affinity chromatography. Using poly(A) polymerase, we add poly(A) tails of over 200 nucleotides to the 3’-OH ends of all linear RNA impurities in a circular RNA preparation. These linear RNAs are then separated from the circular RNA via oligo(dT) affinity chromatography. The linear impurities, bearing longer poly(A) tails, exhibit extended retention on the oligo(dT) column, whereas the circular RNA, with a shorter poly(A) sequence, elutes more quickly. By stepwise elution with ultrapure water, we effectively separate the circular RNA from the linear contaminants. This method offers a fundamental solution to the purification of circular RNA, combining high efficiency, purity, and cost-effectiveness.

## INTRODUCTION

mRNA has recently come into focus as a new drug modality with great therapeutic potential, for the success to addressing the severe acute respiratory syndrome coronavirus 2 (SARS-CoV-2) pandemic^1^. Despite the clear demonstration of efficacy for infectious disease and cancer vaccines, application of mRNA for non-vaccine therapeutics has been limited by the duration of expression, stability and immunogenicity. Various technologies have been developed as soon as possible to weaken its immunogenicity and simultaneously improve mRNA stability and protein expression ability, but their effectiveness still cannot meet the needs of treating non communicable diseases, especially genetic defects^2^. CircRNAss (circRNAs) are distinguished with classic mRNA by their covalently closed loop structure; and the engineering circRNAs have also been found to express protein inside cells^3^.

As a new generation of therapeutic RNAs, circRNAs may solve some of the limitations of classic mRNA including increased stability and reduced immunogenicity^3,4^. At present, various methods for synthesizing circRNAss have been developed, such as chemical synthesis and DNA/RNA ligase synthesis^5^. Among them, the permuted intron–exon (PIE) system with group I/II introns method is the most widely used and can be produced on a large scale^6–9^. However, the process of synthesizing circRNAs from in vitro transcription (IVT) using PIE method generates various RNA impurities, such as dsRNA, precursor RNA, intron RNA, error splicing RNA, and nicked RNA, and the base composition and length of nicked RNA are exactly the same as the circRNAs^3,10,11^. Therefore, the purification of circularized RNA will be a huge challenge. These RNA contaminants can impair the function of circRNA and increase its immunogenicity, greatly affecting the application of circRNAs in biomedicine^8,12^.

There are currently multiple solutions for removing different impurities in the production process of circRNAs. For dsRNA, it can be removed post transcription by RP-HPLC and cellulose fibers-ethanol chromatography^13,14^; it also can be removed during transcription by mutant T7 RNA polymerases and addition of denaturing agents^15–17^. For other types of impurities, there are also some solutions available. For example, RNase R digestion can be used to purify circRNAs^18^. RNase R has the ability to degrade linear RNA, while circRNAs is resistant to its action, allowing RNase R to preferentially hydrolyze the linear RNA in the system while retaining the circRNAs. However, RNase R lacks good specificity, and during the hydrolysis process, a significant amount of circRNAs is also degraded. Additionally, RNA with unique secondary structures may also affect the activity of RNase R^19^. Therefore, this method requires optimization of reaction conditions and often results in low yields^20^. Another method is size exclusion chromatography (SEC) for circRNAs purification^6,21^. This method also has significant drawbacks: first, the amount of RNA purified in a single run is relatively small, making large-scale production difficult; second, while SEC is effective in removing RNA with large differences in molecular size, such as intronic RNA, it is less effective at removing precursor RNA and nicked RNA. IP-RP HPLC was used to purify circRNAs by Kyung Hyun Lee et^8^. This method has significant advantages compared to SEC, firstly, the resolution is improved, and secondly, the purification amount can be linearly amplified at low cost. However, the biggest drawback of this method is the need to introduce toxic organic reagents, such as hexamethylammonium acetate, during the purification process. The residue of toxic reagents is not allowed in the development and production process of biopharmaceuticals.

In this study, we will establish a poly(A) polymerase and oligo(dT)-dependent purification system (PAPdT) for ultra-pure circRNAs to overcome the economic and industrial challenges associated with circRNAs based therapies. First, we utilize a mutant T7 RNA polymerase for the in vitro transcription (IVT) reaction to minimize the introduction of double-stranded RNA (dsRNA) during the initial steps of circRNAs production. Next, we employ Poly(A) polymerase to add a poly(A) tail exceeding 200 nucleotides to the 3’-OH ends of all linear RNAs in the system. Finally, we purify circRNAs from linear RNA using oligo(dT) affinity chromatography. Since the length of the poly(A) tails on linear RNAs is significantly greater than that of the circular RNA itself, these two RNA species can be fractionated and eluted separately, resulting in the isolation of highly pure circular RNA. In addition, we will compare circRNAs purified using the new method with that obtained through traditional methods, assessing differences in protein expression levels and duration of expression in both cellular and animal models.

## MATERIALS AND METHODS

### Expression and purification of Low dsRNA T7 RNA polymerase and Poly (A) polymerase

The T7 RNA polymerase gene, incorporating the G47A and 884G double mutations^16^ and poly (A) polymerase gene^22^ were chemically synthesized by Dynegene Technologies and cloned into the pET28a (+) plasmid (pET28a-T7-GAG and pET28a-PAP1). A 6×His tag was added to the 5’ terminus of the genes to facilitate protein purification via metal chelation affinity chromatography. The pET28a-T7-GAG and pET28a-PAP1 were transformed into E. coli strains BL21(DE3). First, single colony was picked into LB media and allowed to grow 18hr at 37 °C with constant agitation at 200-250 rpm. The seed were then used to inoculate one liter of LB media and grow with constant agitation at 160-200 rpm at 37 °C until OD600 = 0.8. At this point, protein expression was induced by adding IPTG to a final concentration of 0.5 mM and incubated by 16 hr at 160 rpm and 18 °C.

Cells were collected by 4000g centrifugation and resuspended in a lysis buffer (20 mM Tris-HCl,300 mM NaCl,10 mM Imidazole,pH 8 and lysozyme at ∼0.2 mg/mL). Subsequently, cells were incubated at room temperature for 20-30 min with shaking, and the resuspended cells were then disrupted by sonication. Then, the homogenate was centrifuged at 14000g, and the soluble fraction was collected for further steps.

Proteins were purified from the clarified lysate through two sequential affinity chromatography steps^23^. Initially, nickel affinity chromatography was performed, achieving a target protein purity of over 80%. Subsequently, heparin affinity chromatography was employed as a secondary purification step to simultaneously remove protein and nucleic acid contaminants, concentrate the proteins into a smaller volume, and facilitate buffer exchange into appropriate storage buffers (KH2PO4 20mM PH7.5, NaCl 100mM, Glycerol 50%, DTT 10mM, EDTA 0.1mM), and stored at −20°C until further use. The enrichment of purified enzymes in the eluted fractions was assessed by SDS-PAGE.

### Template plasmid construction for circRNA

The DNA template for circRNA synthesis was were chemically synthesized by Dynegene Technologies and cloned to pUC57-AMP plasmid. The template includes the T7 promoter, anabaena 3.0 ribozyme, the internal ribosome entry site IRES-CVB, 5’ and 3’ homologous arms, 5’ and 3’ spacer, and open reading frame (ORF) elements (eGFP or Gaussia Luciferase) and Xba I Enzyme cleavage site^21^. The sequence information of the DNA template is provided in Supplementary Table S1.

### RNA synthesis and circularization

The DNA template was linearized by XbaI digestion and purified with a PCR cleanup kit from Beyotime Biotechnology. The linearized DNAs were used as a template for in vitro transcription with homemade T7 RNA polymerase. The transcription reaction was set up with 1 µg of linearized DNA template, 100 U of homemade T7 RNA polymerase, 100 mM NTPs 2.5μl each, and 10x transcription buffer (Tris-HCl 400mM, Mg(CH_3_COO)_2_ 460mM, spermidine 20mM, DTT 100mM) 2μl, RNase Inhibitor 100U and Inorganic Pyrophosphatase 1U in a total volume of 20 µL^24^. IVT reaction was incubated at 37°C for 3 hours. Following the IVT reaction, 4 U of DNase I (Beyotime Biotechnology) was used to digest the DNA template at 37°C for 15 minutes. And then the product was heated to 70L for 5 min, followed by immediate cooling on ice for 5 min. Subsequently, GTP (Synthgene Biotechnology, Cat# GTP002T-1) and magnesium chloride were added to the RNA to a final concentration of 2 mM and 54mM. The reaction mixture was incubated at 55◦ C for 15 min, after which the RNA was purified by LiCl precipitation.

### Purification of circRNAs by poly(A) polymerase tailing and oligo(dT) affinity chromatography

Adding Poly(A) Tail to 3’ End of All Linear RNA Contaminants: a 1 ml reaction mixture was prepared containing RNA (500 µg), 100 µl of 10x reaction buffer (500 mM Tris-HCl, pH 8.1, 25°C, 2.5 M NaCl, 100 mM MgClL), 20 µl of 100 mM ATP, 1000 U of RNase inhibitor, 250 U of poly(A) polymerase, and DEPC-treated water to a final volume of 1 ml. The reaction was incubated at 37°C for 2–3 hours, and the, 20 µl of 500 mM EDTA was added to terminate the reaction, followed by the LiCl precipitation to concentrate RNA. The poly(A)-tailed RNA was then resuspended in DEPC-treated water for further purification.

Purification of poly(A)-tailed RNA was carried out using a GE AKTA Purifier 100 FPLC system. The system was first cleaned with DEPC-treated water for 10 minutes at a flow rate of 1 ml/min. A 1 ml Proteomix POR50-dT20 prepacked column (Sepax Technologies) was installed and equilibrated by washing with 1x binding buffer (10 mM Tris-HCl, 5 mM EDTA, 1.0 M NaCl, pH 7.0) at 0.5 ml/min. The RNA sample (200 µl, ∼500 µg) was mixed with 200 µl of 2x binding buffer, and 0.8 ml of the mixture was injected into the system at a flow rate of 0.1 ml/min. UV and conductivity detectors were monitored for 30–50 minutes to observe the binding process. Non-specifically bound contaminants were washed off using washing buffer (10 mM Tris-HCl, 5 mM EDTA, 0.2 M NaCl, pH 7.0) for 5 column volumes at a flow rate of 0.5 ml/min. Poly(A)-tailed RNA was eluted with DEPC-treated water at a flow rate of 0.5 ml/min, and two elution peaks were collected. The first peak contained the purified circular RNA, while the second peak corresponded to the poly(A)-tailed linear RNA. The circular RNA fraction was concentrated using an Amicon® Ultra Centrifugal Filter (30 kDa MWCO) and stored at −80°C. This purified RNA, eluted with DEPC-treated water, was ready for subsequent cellular and animal experiments.

### Purification of circRNAs by RNase R

The LiCl-precipitated IVT samples were incubated at 37◦ C for the indicated amounts of time in 20μl reactions that contained 2μl 10× RNase R Buffer (0.2 M Tris–HCl (pH 8.0), 1 mM MgCl2 and 1 M KCl) and the indicated amounts of RNase R. Reactions were purified with LiCl precipitation and the RNA was desolved in 25μl nuclease-free water.

### Purification of circRNAs by SEC column

A GE AKTA Purifier 100 FPLC system was used to purify circRNA. 1mg LiCl-precipitated IVT samples RNA was loaded each run onto a 21×300mm SEC column (Sepax, SRT SEC-1000A, 1000Å pore diameter, 5.0μm particle size, Suzhou, China) with mobile phase containing 10mM Tris, 1mM EDTA, 75mM PB, pH7.4 at 25°C with flow rate 10ml/min. Fractions were collected as indicated and testified with agarose gel electrophoresis.

### Determination of %dsRNA content

The circRNAs were spotted onto positively charged nylon membranes (Nytran SC, Sigma Aldrich). The membranes were then blocked with 2% (w/v) BSA in TBS-T buffer (20mM Tris–HCl, 150mM NaCl, 0.1% Tween-20, pH 7.4), and incubated with anti-dsRNA monoclonal antibody J2 (Jena Bioscience) at 4°C overnight. Membranes were washed with TBS-T buffer 3 times and incubated with HRP-conjugated goat anti-mouse IgG (Beyotime Biotechnology). Following additional 3x wash with TBS-T, the membranes were treated with ECL Plus Western blot detection reagent (Beyotime Biotechnology). Images were captured using a digital imaging system.

### Determination of circRNA by by HPLC-SEC

HPLC-SEC experiments were performed using a Shimadzu LC20 HPLC system equipped UV detector with a 4.6×300mm SEC column (Sepax, SRT SEC-1000A, 1000Å pore diameter, 5.0μm particle size, Suzhou, China). 3ug circRNA was loaded each run onto the column with mobile phase containing 10mM Tris, 1mM EDTA, 20mM PB, pH7.4. The SEC separation was performed at a flow rate of 0.35 mL/min at 25 °C; the UV detection wavelength was 260 nm.

### Measurement of the translation products from circRNAs

Cells were seeded into 24-well plates one day before transfection. The purified circRNAs are transfected into cells using Lipo6000 (Beyotime Biotechnology) according to the manufacturer’s manual. After transfection, cells were cultured at 37°C for 24h. the eGFP green fluorescent protein was observed and photographed using Olympus fluorescence microscope IX73.

### Real-time quantitative PCR

samples from different groups were used for gene expression analysis by an RT-PCR. Amplification and detection using TIANGEN Talent qPCR PreMix were performed with an ABI Prism 7500 sequence detection system. All the primers used in this study have been listed in Supplemental Table 1. All the expression levels of the target genes were normalized to the expression of GAPDH.

### Lipid nanoparticle (LNP) formulation and characterization

The synthesis of lipid nanoparticles (LNPs) was conducted using a microfluidic approach. A lipid solution in ethanol was prepared containing SM-102 (Hefei HuaNa Biomedical Technology), DSPC (Hefei HuaNa Biomedical Technology), cholesterol (AVT), and DMG-PEG2000 (Hefei HuaNa Biomedical Technology) in a molar ratio of 50:10:38.5:1.5. This lipid mixture was subsequently combined with an aqueous phase containing circRNAs, which were diluted in a 10 mM citrate buffer (pH 4), to achieve an ionizable lipid/RNA weight ratio of 10:1. LNP formulation was carried out using a Nano Plus microfluidic device (Nexstarbio, Shanghai), set to a total flow rate of 10 mL/min, with an aqueous-to-ethanol volume ratio of 3:1. Following the formation of the LNPs, ethanol was removed via dialysis using an Amicon® Ultra Centrifugal Filter (30 kDa MWCO).

For LNP characterization, the average particle size, polydispersity index (PDI), and zeta potential were measured using a Malvern Zetasizer Advance Lab Blue Label (Malvern Instruments Ltd). The encapsulated RNA content was quantified using the Quant-IT® Ribogreen assay (Invitrogen) according to the manufacturer’s protocol, and LNP encapsulation efficiency was confirmed by agarose gel electrophoresis. The final LNP formulation was adjusted to an mRNA concentration of 100 μg/mL and stored at 4°C for subsequent in vivo studies.

### Administration of LNP-circRNAs in mice

Male C57BL/6J mice (7–8 weeks old) were procured from Charles River Laboratories (Beijing, China). The animals were housed under standard conditions with a 12-hour light/12-hour dark cycle, a relative humidity of 60 ± 5%, and a temperature of 25 ± 1 °C, with ad libitum access to food and water. All procedures were performed in accordance with the ethical guidelines approved by the Ethics Committee of Zhengzhou University.

A total of 20 µg of LNP-circRNAs in PBS were administered intravenously into the mice. Blood samples (10 µL) were collected daily from the tail vein using 50 mM EDTA as an anticoagulant over a two-week period. For measurement of Gluc activity, a 100 mM coelenterazine solution was prepared by diluting it in PBS containing 5 mM NaCl (pH 7.2). A 5 μL aliquot of blood was transferred to a well of a 96-well black plate. Gluc activity was quantified using a plate luminometer, which was programmed to inject 100 μL of 100 mM coelenterazine and measure photon counts for 10 seconds. Data analysis was performed by plotting the relative light units (RLU; y-axis) as a function of time (x-axis).

### Statistical analysis

Data were expressed as mean ± standard error of the mean (SEM). Statistical significance was assessed using either a two-tailed unpaired Student’s t-test for comparisons between two groups. A p-value less than 0.05 was considered statistically significant. All statistical analyses were conducted using Prism version 8.0.

## RESULTs

### 1. Production of Low dsRNA T7 RNA polymerase mutant and Poly (A) polymerase

A double-mutant T7 RNA polymerase (T7-GAG, G47AL+L884G) has been reported to significantly reduce double-stranded RNA (dsRNA) byproducts during in vitro transcription (IVT) while maintaining RNA yield and purity^16^. In this study, we synthesized and expressed T7-GAG based on published protocols and validated its IVT performance. Additionally, E. coli Poly(A) polymerase (PAP) was cloned and expressed following established methods^25^. The genes encoding T7-GAG and PAP were synthesized and inserted into the pET28a vector with an N-terminal His tag for purification. Figures 1A and 1B show the expression and purification of T7-GAG and PAP enzymes in E. coli. After two-step purification via nickel affinity and heparin affinity chromatography, both enzymes were obtained with >90% purity.

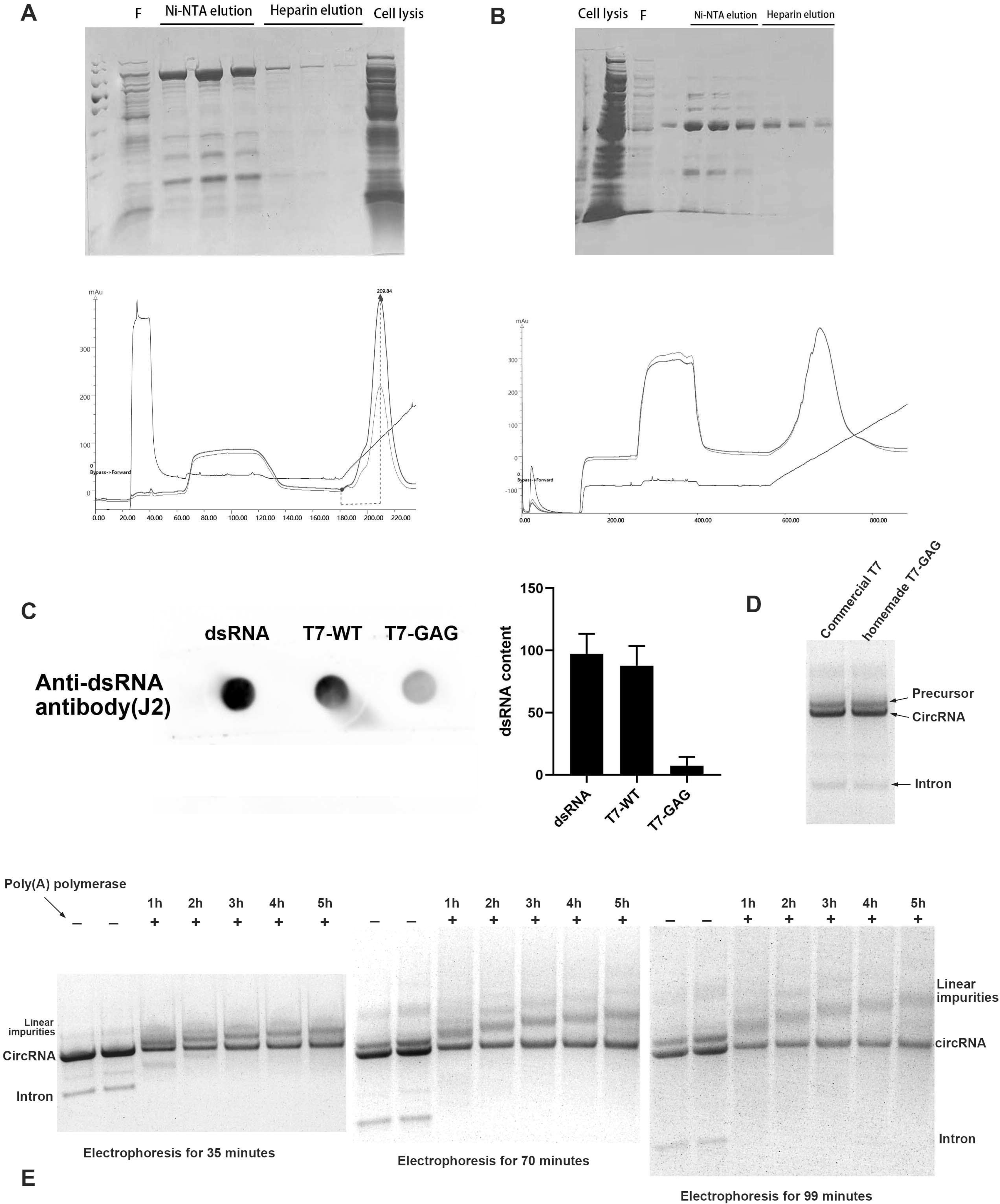

IVT was performed using purified T7-GAG RNA polymerase, and dsRNA byproducts were quantified using the J2 antibody. The results showed that dsRNA yield from T7-GAG was significantly lower than that of commercial wild-type T7 RNA polymerase (Figure 1C), while total RNA yield was similar between the two enzymes (Figure 1D). In the crude product of circularized RNA, only the circular RNA cannot be polyadenylated, while the 3’ ends of other linear impurities were polyadenylated to varying lengths under the E. coli Poly(A) polymerase catalytic reaction. As shown in Figure 1E, poly(A) tailing of all RNAs was achieved after 1-5 hours, as confirmed by electrophoresis at intervals from 35 to 99 minutes. The electrophoresis results showed gradual separation of linear and circular RNA, with circular RNA remaining in place.

### 2. The oligo(dT) column effectively distinguishes between circular RNA and all the linear RNA with poly(A) tails

Based on the above results, we successfully obtained two purified proteins with expected enzyme activity and functionality. These enzymes were used to produce circular RNA, which was then purified using an oligo(dT) chromatography column. We synthesized and ligated several genetic elements, including the eGFP gene, Anabaena type I intron, IRES-CVB3 sequence, spacer sequence, and homologous 5’ and 3’ sequences, into the pUC57 Kan vector. The plasmid was amplified in *E. coli*, linearized, and in vitro transcribed (IVT) using the T7-GAG enzyme to generate eGFP circular RNA precursors. Cyclization was induced by adding GTP and Mg^2 +^, as shown in Figure 1D. During IVT, some RNA underwent autonomous cyclization, and after the cyclization step, approximately 90% of the RNA was cyclized. The crude circular RNA was purified using lithium chloride precipitation, followed by poly(A) tailing for 2 hours with poly(A) polymerase, and the reaction was terminated with EDTA.

For purification, 500 µg of polyA-tailed circular RNA was diluted with 2X binding buffer and applied to an oligo(dT) column (Figure 2A). The chromatogram shows that both circular RNA and linear RNA impurities bind to the column due to the presence of a 50-bp poly(AC) sequence in the circular RNA. Linear RNA impurities, which have longer poly(A) tails, exhibit longer retention times during washing with DEPC-treated water, while circular RNA elutes more quickly. The two elution peaks were concentrated and analyzed by agarose gel electrophoresis. As shown in Figure 2B, the first elution peak contained a single band corresponding to purified circular RNA, while the second peak showed multiple bands, indicating a mixture of linear RNA impurities and a small fraction of circular RNA. To assess the effect of poly(A) tail length on purification, we varied the poly(A) tailing time to produce impurities with different tail lengths, then purified them by oligo(dT) chromatography. As shown in Figure 2C, the length of the poly(A) tail did not affect the purification efficiency.

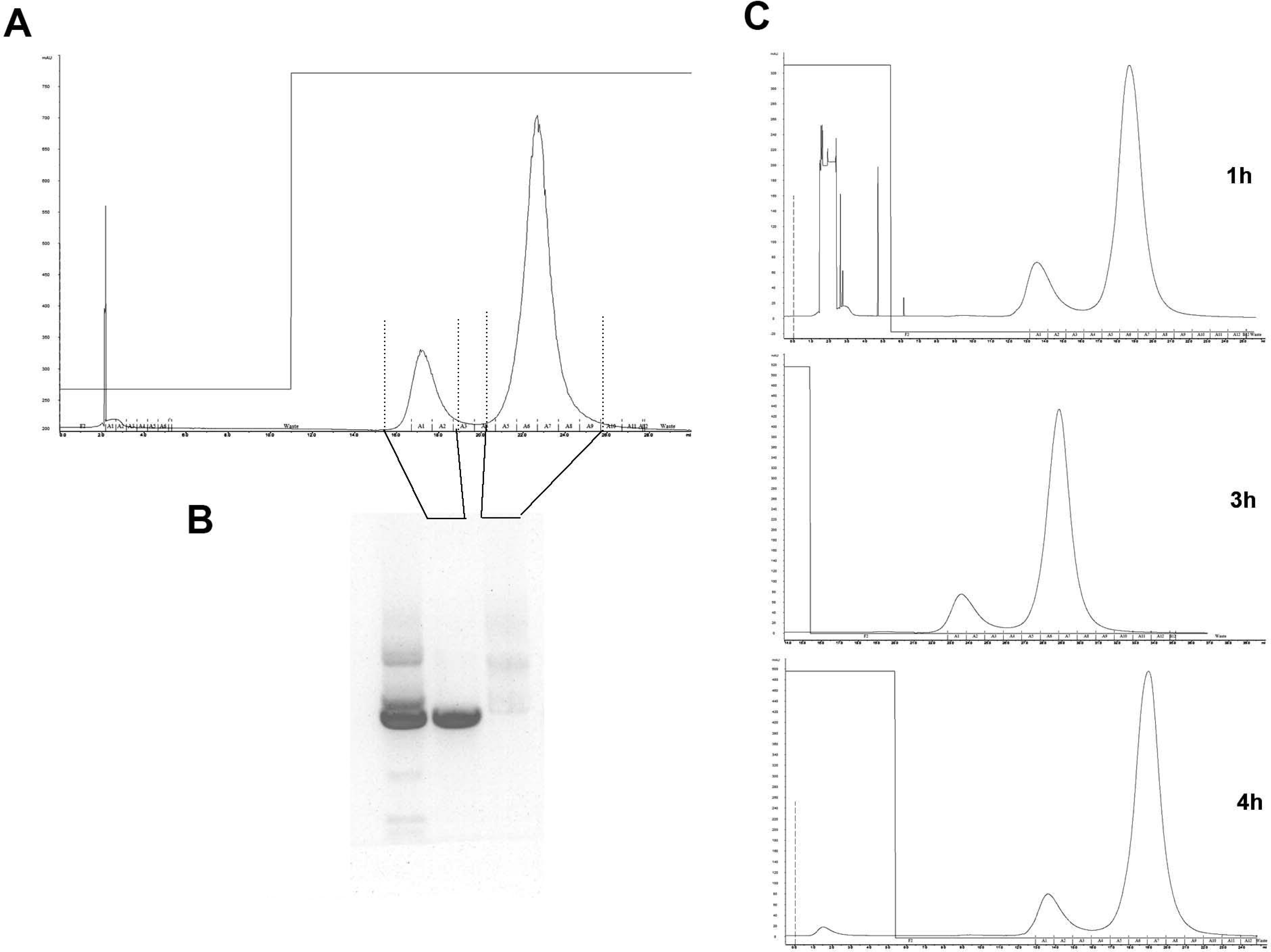

### 3. RNase R and HPLC-SEC purification methods cannot obtain a single circular RNA band

We compared the new purification method proposed in this study with traditional RNase R treatment and HPLC-SEC methods for isolating eGFP circular RNA. RNase R selectively digests RNA with 3’-OH termini, retaining circular RNA, but its specificity is limited, and it can also degrade circular RNA, although to a lesser extent. To achieve optimal purification, digestion conditions must be optimized for each new circular RNA, as shown on the left side of Figure 3A. Under moderate digestion conditions, we obtained relatively pure circular RNA, though some RNA was lost in the process, as shown on the right side of Figure 3A. This is consistent with previous studies^3,10,11^.

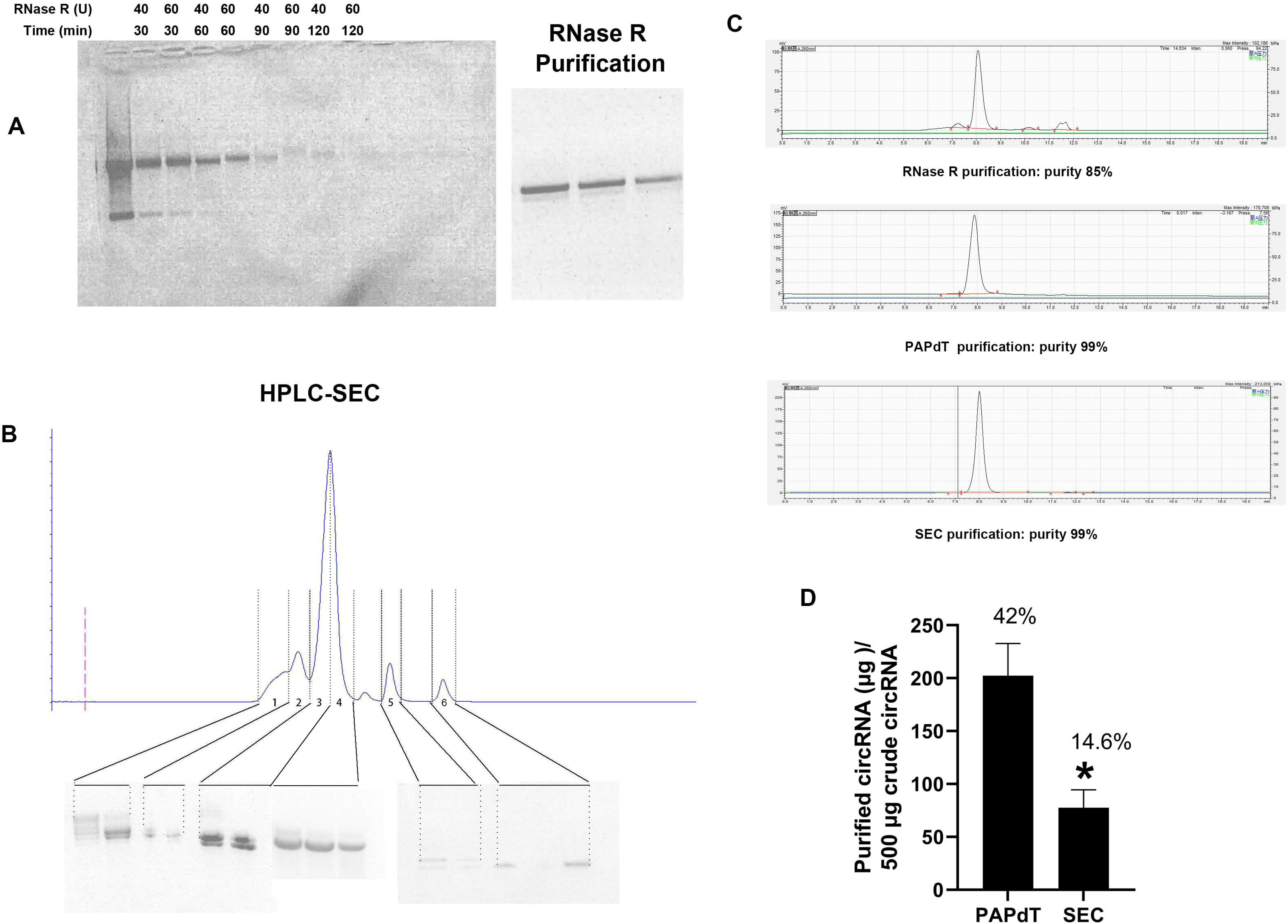

The HPLC-SEC method separates components by molecular size (Figure 3B). The chromatogram shows multiple peaks corresponding to different components in the circular RNA preparation. Partial collection was performed based on the elution peaks. Agarose gel electrophoresis of each elution fraction revealed that circular RNA was mainly concentrated in the third and fourth elution peak. As shown in the figure, the samples at positions 3 and 4 had the highest content of circular RNA, of which circular RNA in position 3 accounted for about 50%. There were three tubes of samples collected at position 4, with a small amount of impurities in the first tube, and no impurities were seen in the second and third tubes on the electrophoresis graph. Subsequent experiments mainly used samples from the second and third tubes at position 4.

To further evaluate the purity of circular RNA obtained from the three purification methods, we conducted an analysis using HPLC-SEC, as presented in Figure 3C. The RNA purified via RNase R exhibited four distinct elution peaks, corresponding to a purity of approximately 85%. In contrast, the RNA purified by SEC displayed one peak, with a purity exceeding 99%. The RNA purified using the PAPdT also exhibited a single peak, with purity greater than 99%. When comparing the recovery rates of the two purification methods, we found that SEC yielded approximately 74 μg of ultra-pure circular RNA from 500 μg of circular RNA stock, whereas the PAPdT method produced over 200 μg of ultra-pure circular RNA. Thus, the purification efficiency of PAPdT was calculated to be 42%, while that of SEC was only 14.6%.

### 4. The circular RNA obtained by PAPdT has higher protein expression levels and longer duration in cells and animals

We then compared the protein expression levels of circRNA obtained by the PAPdT and SEC purification method. As shown in Figure 4A, 3 μg of eGFP circular RNA obtained by the two purification methods were transfected into BV-2 cells, and the cells were observed and compared using a fluorescence microscope every 24 hours. The results showed that 24 hours after transfection, the expression of eGFP protein in RNA purified by PAPdT far exceeded that of RNA purified by the SEC method. The protein expression of RNA purified by the SEC method decreased significantly after 72 hours, while the expression of RNA purified by PAPdT was slightly reduced; even after one cell passage, the RNA purified by the PAPdT still had significant protein expression.

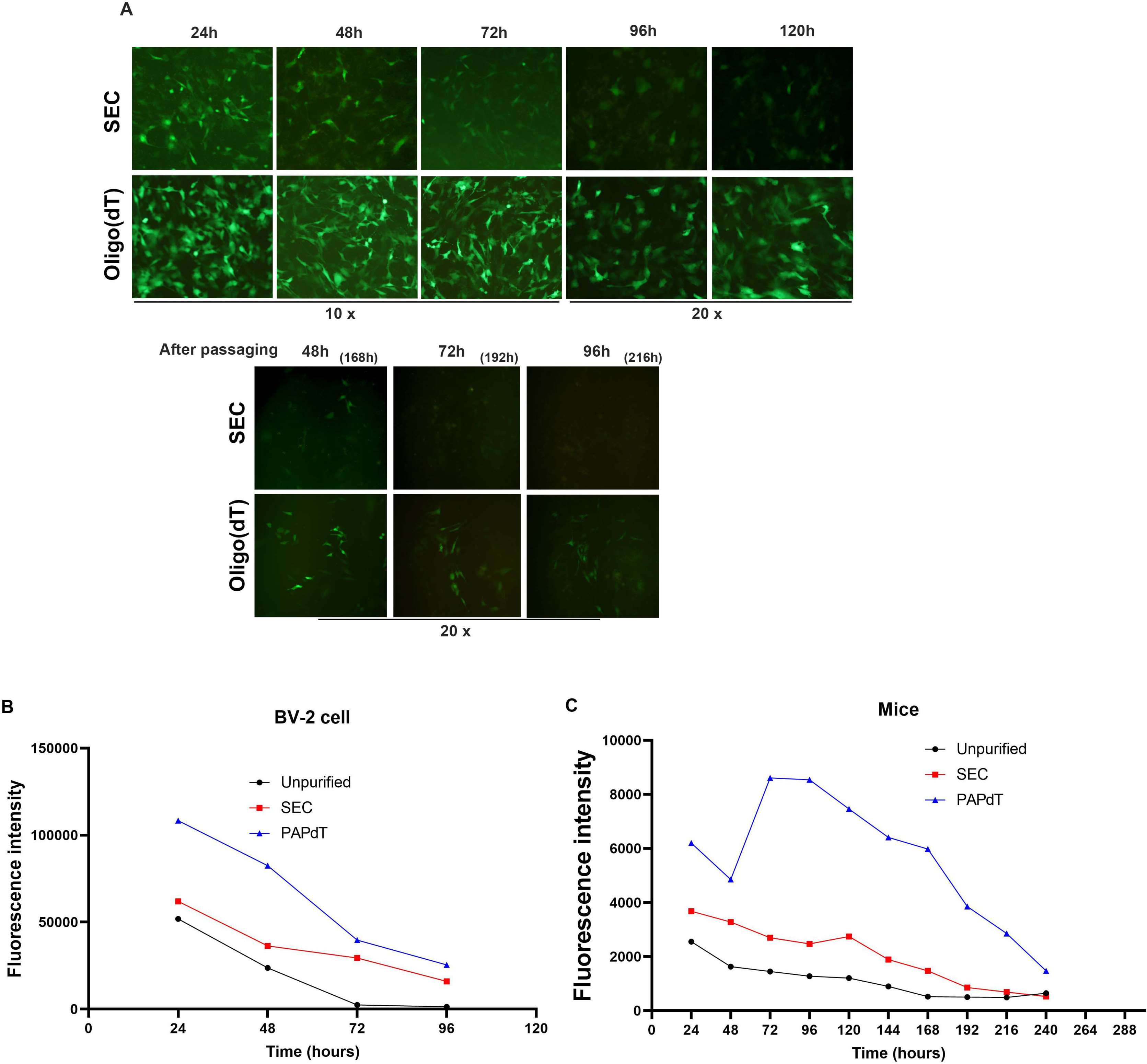

To further verify the effectiveness of the PAPdT purification method, we produced extracellular secretory hGluc circRNA and purified it using both PAPdT and SEC methods. Finally, we produced lipid nanoparticles (LNPs) encapsulating the circRNA and detected the level of protein expression in cells and animals. As shown in Figure B, the fluorescence value of RNA purified by PAPdT was twice that of the SEC purification method after the first 24 hours, and its high level of expression lasted for more than 4 days. Figure 4C shows the expression level of hGluc in mouse serum. It can be seen that the fluorescence value of RNA purified by PAPdT is much higher than that of the SEC purification method, and it shows a longer expression capacity.

### 5. The circular RNA obtained by the new method has extremely low immunogenicity

The immunogenicity of circular RNA is closely related to the purity of RNA, especially dsRNA is an important impurity that provokes immune response. In this project, we eliminated the introduction of dsRNA from the first step of IVT process, and then obtained ultra-pure circRNA with a purity of more than 99% through our PAPdT purification step. This RNA should have very low immunogenicity. To verify this point of view, we used LNP to encapsulate and transfect A549 cells with unpurified RNA obtained by WT T7 RNA polymerase, SEC-purified RNA, and PAPdT-purified RNA, and used fluorescent quantitative PCR to detect the expression of inflammatory factors. As shown in Figure 5, unpurified RNA provokes significant expression of immune factors, while SEC-purified and PAPdT-purified RNA hardly provokes immune response.

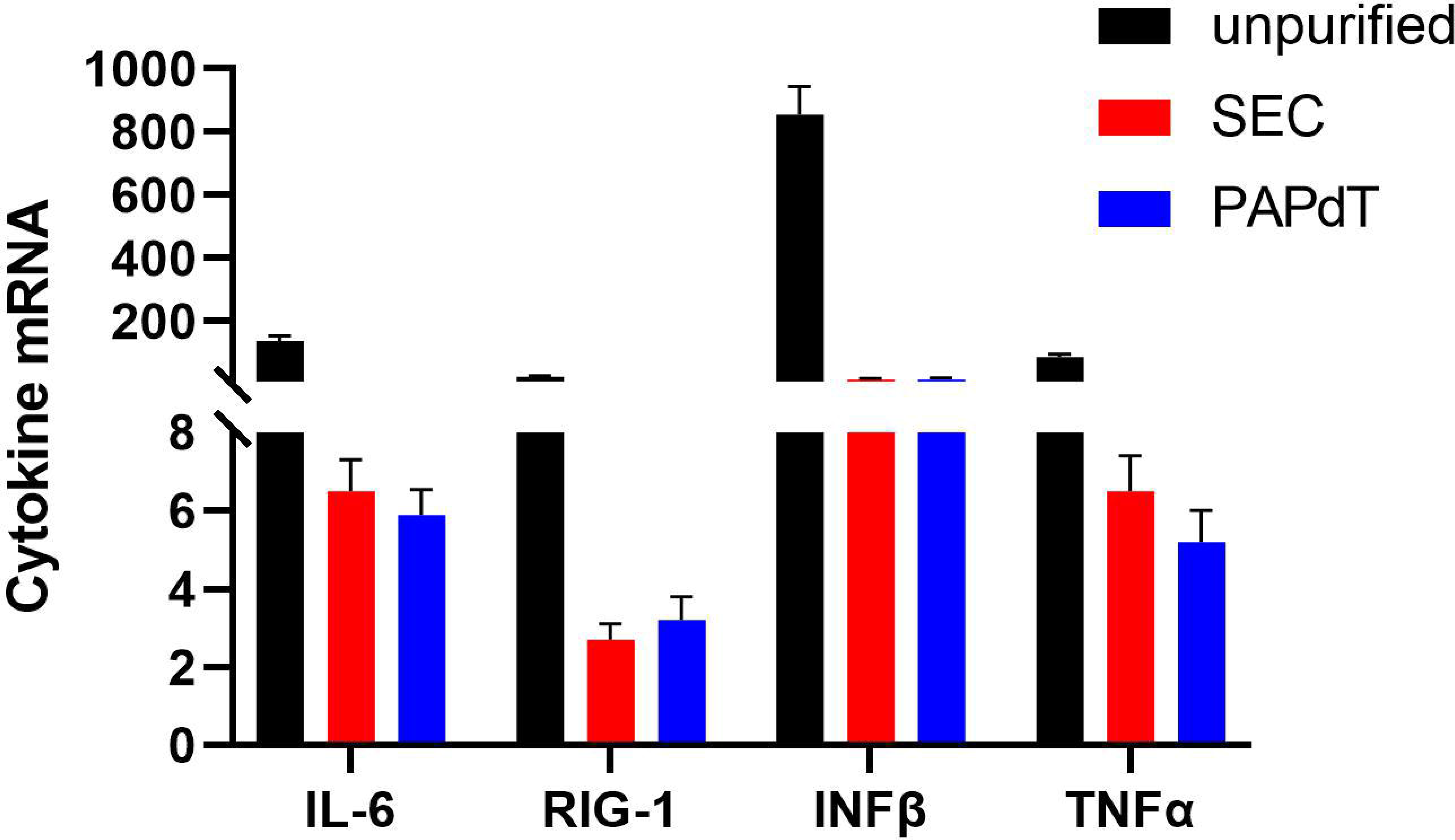

## DISCUSSION

Since the pioneering discovery that engineered circular RNA (circRNA) can express target proteins and hold potential for medical applications, research on its functions has advanced rapidly^3^. Various strategies for generating circRNA have been developed, and many biopharmaceutical companies are adopting these methods^10,26,27^. However, the large-scale production of circRNA faces challenges due to impurities such as cut introns, precursor RNA, open-circular RNA, spliced RNA errors, and double-stranded RNA (dsRNA). Consequently, the purification of these impurities remains a key obstacle to the widespread production and application of circRNA. Several purification techniques have been developed to address this issue. For dsRNA removal, using a mutant T7 RNA polymerase to prevent its introduction during in vitro transcription (IVT) is a cost-effective and reliable approach^27^.

One of the earliest purification methods for circRNA is RNase R digestion, including its modified version, RPAD^28,29^. This technique utilizes RNase R, which selectively degrades linear RNA with 3’-OH ends, while closed circular RNA remains intact. However, some linear RNAs also show resistance to RNase R, particularly when they adopt specific secondary structures. To overcome this, poly(A) polymerase tailing is used, followed by RNase R digestion, as seen in the RPAD method. Despite its utility, this approach does not achieve complete purification of circRNA and requires extensive optimization of digestion time and enzyme-to-RNA ratios. Moreover, excessive RNase R can degrade circRNA, and the method is not scalable for large RNA batches. Consequently, it is mainly used for circular RNA enrichment in sequencing applications.

Another widely employed purification method is high-performance liquid chromatography (HPLC), specifically size-exclusion chromatography (HPLC-SEC) or ion-pair reverse-phase HPLC (IP-RP-HPLC). HPLC-SEC is effective for achieving high-purity circRNA, and several pharmaceutical companies have adopted this technique. However, the major limitations include the high cost of the purification column and the small amount of RNA that can be processed per run. For instance, a typical SEC column like the SRT SEC-1000 30×300mm from Sepax Technologies can process only up to 6 mg of RNA, with a price exceeding 120,000 RMB. Therefore, while HPLC-SEC is suitable for preclinical research, it is cost-prohibitive for industrial-scale production. IP-RP-HPLC addresses some of these issues by improving the yield, but the use of toxic reagents limits its application in biopharmaceutical settings.

Recently, two new purification methods have been introduced: affinity chromatography using oligo(dT) and ultrafiltration^30,31^. However, these methods have significant drawbacks, such as low purity. In the work by Diego Casabona et al., affinity chromatography using a 50-bp poly(AC) sequence and an oligo(dT) column was ineffective due to the presence of similar sequences in precursor RNA, open-circular RNAs, and spliced RNAs, preventing successful purification. Similarly, the ultrafiltration method developed by Scott M. Husson et al. achieved only 86% purity for circRNA. Thus, no single purification method has yet managed to balance purity, yield, and cost effectively.

In contrast, our PAPdT method offers a solution that addresses these challenges. In this approach, poly(A) polymerase is used to add long poly(A) tails to all linear RNA impurities in the circRNA preparation. These poly(A)-tailed impurities bind more strongly to the oligo(dT) column, allowing the circular RNA and linear impurities to be efficiently separated during affinity chromatography. This method achieves exceptionally high yield and purity. For example, from 500 μg of stock solution, over 200 μg of ultra-pure circRNA can be obtained, whereas the HPLC-SEC method typically yields only 70 μg of pure circRNA (figure 3D). Moreover, the cost of the PAPdT method is low, as it utilizes commonly available oligo(dT) columns and poly(A) polymerase, an enzyme that is easy to express, purify, and is highly stable. The mild reaction conditions and high compatibility of poly(A) polymerase make this method highly suitable for industrial-scale production without the need for extensive optimization.

Furthermore, our method is versatile, applicable to circRNA produced by other techniques, and can be adapted for circRNA that lacks a poly(A) sequence. In such cases, the circular RNA will flow through the column, while the linear impurities are retained. This ensures that higher yields and purities of circRNA can be obtained in these scenarios as well. Our PAPdT method is the first to fully resolve the purification challenges in circRNA development and production, making it a promising solution for both research and industrial applications. In addition, our method yielded results consistent with those of Diego Casabona et al. The circRNA isolated via oligo(dT) affinity chromatography demonstrates higher expression levels and extended stability compared to circRNA obtained through alternative methods. Notably, if a mutant T7 RNA polymerase is employed during the in IVT step to fully prevent the introduction of dsRNA, this approach ultimately produces ultra-pure circular RNA, suitable for direct application in subsequent preclinical and clinical studies.

## Author Contributions

Conceptualization: Lulu Wei, Yingwu Mei; Methodology: Lulu Wei, Mengjuan Zhang, Yingwu Mei; Software: Yingwu Mei; Validation: Lulu Wei, Mengjuan Zhang, Haiyan Zhao; Formal analysis: Haifeng Zhang; Investigation: Lulu Wei, Mengjuan Zhang, Haiyan Zhao, Yingwu Mei; Resources: Haifeng Zhang; Data Curation: Lulu Wei, Yingwu Mei; Writing - Original Draft: Yingwu Mei; Writing - Review & Editing: Lulu Wei, Yingwu Mei; Visualization: Yingwu Mei; Supervision: Yingwu Mei; Project administration: Yingwu Mei; Funding acquisition: Yingwu Mei.

## Funding

This research was funded by the Henan Province Scientific and Technological Research Projects (242102310423).

## Competing Interest Statement

Zhengzhou University has filed a patent application related to the technology described in this paper.

